# MeSH-informed enrichment analysis and MeSH-guided semantic similarity among functional terms and gene products in chicken

**DOI:** 10.1101/034975

**Authors:** Gota Morota, Timothy M Beissinger, Francisco Peñagaricano

## Abstract

Biomedical vocabularies and ontologies aid in recapitulating biological knowledge. The annotation of gene products is mainly accelerated by Gene Ontology (GO) and more recently by Medical Subject Headings (MeSH). Here we report a suite of MeSH packages for chicken in Bioconductor and illustrate some features of different MeSH-based analyses, including MeSH-informed enrichment analysis and MeSH-guided semantic similarity among terms and gene products, using two lists of chicken genes available in public repositories. The two published datasets that were employed represent (i) differentially expressed genes and (ii) candidate genes under selective sweep or epistatic selection. The comparison of MeSH with GO overrepresentation analyses suggested not only that MeSH supports the findings obtained from GO analysis but also that MeSH is able to further enrich the representation of biological knowledge and often provide more interpretable results. Based on the hierarchical structures of MeSH and GO, we computed semantic similarities among vocabularies as well as semantic similarities among selected genes. These yielded the similarity levels between significant functional terms, and the annotation of each gene yielded the measures of gene similarity. Our findings show the benefits of using MeSH as an alternative choice of annotation in order to draw biological inferences from a list of genes of interest. We argue that the use of MeSH in conjunction with GO will be instrumental in facilitating the understanding of the genetic basis of complex traits.

## Introduction

Understanding the genetic basis of variation for complex traits remains a fundamental goal of biology. Different approaches, including whole-genome scans and genome-wide expression studies, have been used in order to identify individual genes underlying economically relevant traits in a wide spectrum of agricultural species. These studies usually generate lists of genes potentially involved in the phenotypes under study. The challenge is to translate these lists of candidates genes into a better understanding of the biological phenomena involved. It is increasingly accepted that overrepresentation or enrichment analysis (Drăghici et al., 2003) can provide further insights into the biological pathways and processes affecting complex traits.

Recently, the Medical Subject Headings (MeSH) vocabulary (Nelson et al., 2004) has been proposed for defining functional sets of genes in the context of enrichment analysis. MeSH is a controlled life and medical sciences vocabulary maintained by the National Library of Medicine to index documents in the MEDLINE database. Each bibliographic reference in the MEDLINE database is associated with a set of MeSH terms that describe the content of the publication. Importantly, MeSH contains a substantially more diverse and extensive range of categories than that of Gene Ontology (GO) (Ashburner et al., 2000), which is probably the most popular among the initiatives for defining functional classes of genes (Nakazato et al., 2008). Therein, GO terms are classified into three domains: biological processes, molecular functions, and cellular components. This ontology has been successfully used for dissecting relevant traits in livestock species (e.g, Peñnagaricano et al., 2013; Gambra et al., 2013). Similarly, each MeSH term is clustered into 19 different categories; some MeSH categories, such as Diseases, are not included in GO, whereas other functional categories, such as Phenomena and Processes or Chemicals and Drugs, share similar concepts with those of GO. The recent availability of MeSH software packages has rendered agricultural species amenable to MeSH-based analysis (Tsuyuzaki et al., 2015). For instance, MeSH enrichment analysis has been successfully applied to mammals including dairy cattle, swine, and horse (Morota et al., 2015), and to maize (Beissinger and Morota, 2016). These studies showed the potential of MeSH for enhancing the biological interpretation of sets of genes in agricultural organisms.

The main objective of the current study was to report the availability of MeSH Bioconductor packages for chicken, and to illustrate the features of different MeSH-based analyses, including MeSH-informed enrichment analysis and MeSH-guided semantic similarity among terms and gene products. For this purpose, we used two lists of selected genes available in public repositories: (i) differentially expressed genes reported in a RNA-seq study (Zhuo et al., 2015) and (ii) candidate genes historically impacted by selection detected in a whole-genome scan using a broad spectrum of populations (Beissinger et al., 2015). The results of the MeSH-based enrichment analysis were contrasted with GO terms. The use of MeSH and GO terms in functional genomics studies can be further explored through computing the similarity between significant functional terms as well as the similarity between significant genes by leveraging the hierarchies of these two controlled vocabularies.

## Materials and Methods

We used two datasets from previously published studies with the objective of demonstrate some capabilities of different MeSH-based analyses in chicken. The first dataset includes 263 genes that showed differential expression in abdominal fat tissue between high and low feed efficiency broiler chickens (Zhuo et al., 2015). The second dataset contains 352 genes identified by a whole-genome scan using Ohta’s between-population linkage disequilibrium measure, 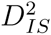, in a panel that included 72 different chicken breeds (Beissinger et al., 2015). In both datasets, the list of background genes was defined as all annotated genes in the chicken genome available in NCBI. Below we present the MeSH analyses coupled with several example code for illustration purposes.

The suite of MeSH (Tsuyuzaki et al., 2015) and the GOstats (Falcon and Gentleman, 2007) packages in Bioconductor were used for performing a hypergeometric test in the enrichment analysis. This test evaluates whether a given functional term or vocabulary is enriched or overrepresented with selected genes. In particular, the *P*-value of observing *g* significant genes in a functional term (i.e. MeSH or GO term) was calculated by

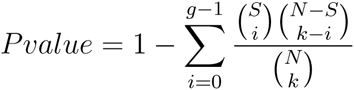

where *S* is the total number of selected genes, *N* is the total number of analyzed genes, and *k* is the total number of genes in the functional term under study. The meshr package has a feature to perform a multiple testing correction by choosing from Benjamini-Hochberg, Q-value or empirical Bayes method. We used a lenient *P*-value 0.05 for the illustrative data in order to directly compare the results from MeSH enrichment analysis with the ones from the GOstats package, which does not offer a multiple testing correction option. Although a multiple testing correction reduces false positives, if we view MeSH analysis as a tool to generate hypotheses or to obtain a big picture of selected genes for subsequent downstream analysis, we may want to know the top 10% of MeSH terms regardless of *P*-values.

The first step of MeSH analysis is to load the namespace of the packages.

`library`(MeSH. db)

`library`(MeSH.Gga. eg. db)

`library`(meshr)

The MeSH.db package contains the relationship between MeSH IDs and MeSH terms. The MeSH.Gga.eg.db is an annotation package that provides the correspondence between MeSH IDs and Entrez Gene IDs. This package was created based on gene2pubmed (ftp://ftp.ncbi.nih.gov/gene/DATA/) that maps Entrez Gene IDs and PubMed IDs. By using data licenced by PubMed (http://www.nlm.nih.gov/databases/license/license.html), we then associated PubMed IDs to MeSH terms. This was followed by merging MeSH terms with MeSH IDs via NLM MeSH (Tsuyuzaki et al., 2015). The meshr package performs a hypergeometric test and returns significantly enriched MeSH terms. Once the three packages are loaded, we proceed to create the object of a parameter class MeSHHyperGParams-class. This object contains all parameters required to run the hypergeometric test.

meshParams <– **new**(”MeSHHyperGParams”, gene 122 Ids = select edGene s, unive r s eGene Ids = univer seGenes, annotat ion = “MeSH.Gga. eg. db”, category = “D”, database = “gene2pubmed”, pvalueCutof f = 0. 0 5, pAdjust = ”none”)

Here geneIds and universeGeneIds are the vectors of Entrez Gene IDs for selected and background genes, respectively, category is one of the abbreviation codes for MeSH categories such as D (Chemicals and Drugs), C (Diseases), A (Anatomy), and G (Phenomena and Processes), pvalueCutoff is the numeric value for *P*-value cutoff, and pAdjust allows users to choose multiple testing methods from BH (Benjamini-Hochberg), QV (Q-value), lFDR (empirical Bayes), and none (unadjusted). Finally, the meshHyperGTest function accepts the MeSHHyperGParams-class object and perform a MeSH enrichment analysis.

`meshR <– meshHyperGTest (meshParams)`

The returned object is MeSHHyperGResult-class and we can access the results with the summary function.

**summary**(meshR)

The summary function returns a data.frame object with information about MeSH ID, *P*-value, MeSH term, Entrez Gene ID, and PubMed ID.

In addition, the hierarchical structures of MeSH and GO permitted us to compute semantic similarities between functional terms (Lord et al., 2003; Pesquita et al., 2009). This is a metric between two terms on the basis of their biological meanings of annotation: the closer two terms are in the hierarchy, the higher the similarity measure is between these terms. Figure 1 shows a MeSH hierarchy for illustrative purpose. In this example, the semantic similarity measure between Mesh Term 2 and Mesh Term 3 is greater than that of Mesh Term 1 and Mesh Term 2 because they are closer in the hierarchy. We employed the information content-based Jiang and Conrath's measure (Jiang and Conrath, 1998) to compute the pairwise similarities within GO ontologies and MeSH headings. The semantic similarity measure between two terms *t*_1_ and *t*_2_ is given by the information content *IC*(*t*) = – log*p*(*t*), where *p*(*t*) is the probability of occurrence of the term *t* and its children terms in MeSH or GO hierarchy. The semantic distance metric is a function of

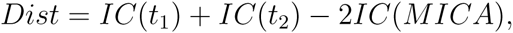

where MICA is the most informative common ancestor.

**Figure 1:**
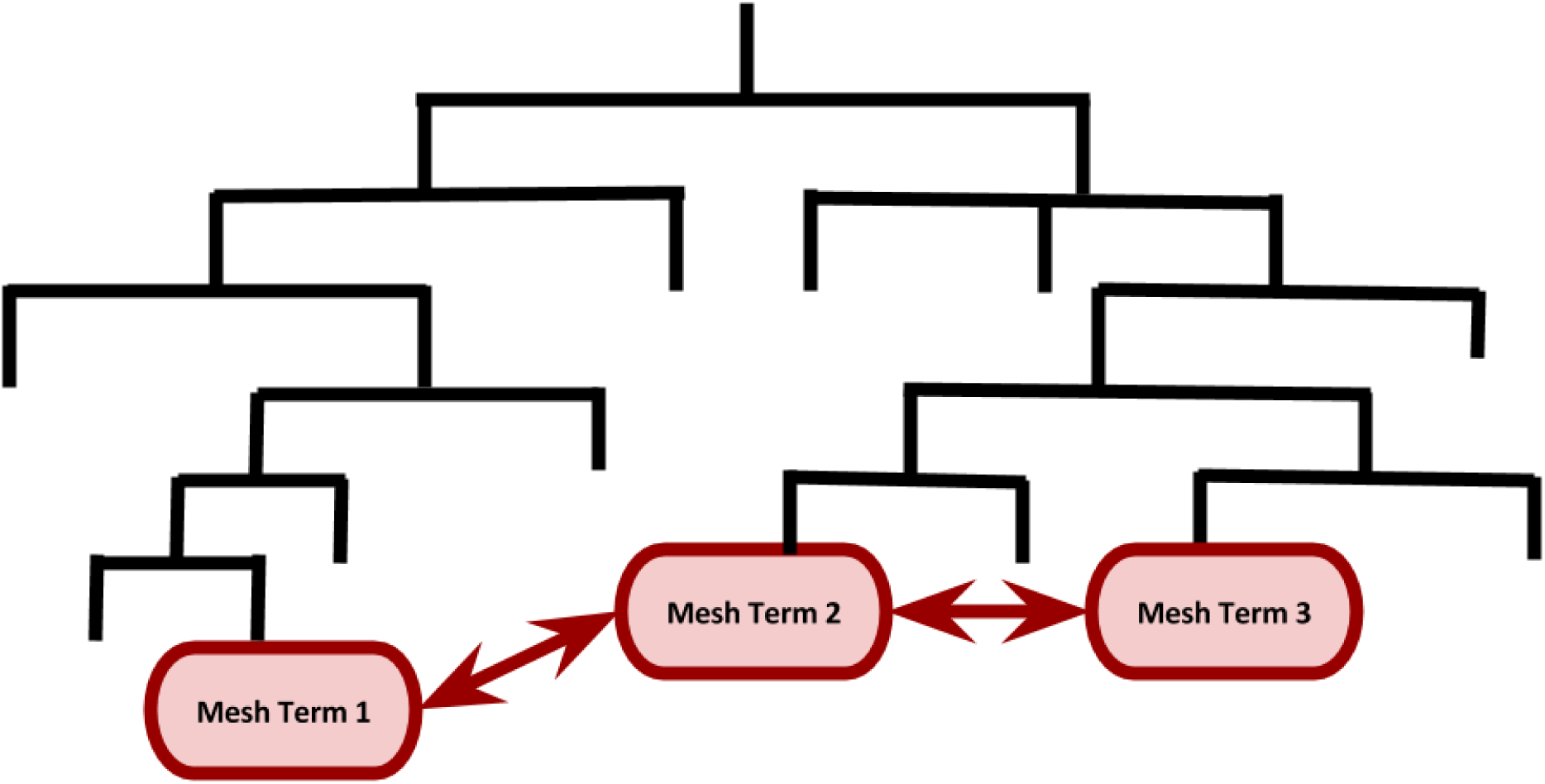
A cartoon illustrating semantic similarity among MeSH terms in the MeSH hierarchy. The semantic similarity measure between Mesh Term 2 and Mesh Term 3 is greater than that of Mesh Term 1 and Mesh Term 2 because they are closer in the hierarchy.

We further computed semantic similarity between selected genes by aggregating their MeSH or GO terms assigned. This is a similarity measure at the level of genes which is analogous to a similarity matrix among SNPs (Morota and Gianola, 2013). We calculated similarity scores over all pairs of terms between the two vocabulary sets of genes under consideration. All these GO and MeSH-guided semantic similarity analyses were carried out using the GOSemSim (Yu et al., 2010) and the MeSHSim (Zhou et al., 2015) Bioconductor packages, respectively. We selected exactly the same genes as were identified in GO categories when computing MeSH-based gene similarity to allow direct comparisons between these two functional vocabularies. Source code and reproducible output reports generated by R Markdown are available as Supporting Files.

## Data Availability

The MeSH.db, MeSH.Gga.eg.db, and meshr packages are availalbe for download at Bioconductor https://www.bioconductor.org/. The two datasets used in the current study have already been published. The gene expression data can be downloaded from http://journals.plos.org/plosone/article?id=10.1371/journal.pone.0135810#sec025. Raw data for the selective sweep data are available from http://dx.doi.org/10.6084/m9.figshare.1497961, and selected genes can be found in Beissinger et al. (2015).

## Results

### Summary of MeSH and GO annotations

The organism and the biomaRt Bioconductor packages were queried to annotate genes by MeSH and GO terms. Table 1 shows the total number of genes (background and selected genes) annotated by MeSH and GO in each of the datasets under study. Both MeSH and GO terms had a similar number of annotated known genes (10,227 vs. 12,460), whereas the number of selected genes with MeSH terms assigned was about one-half of that of GO. For example, in the gene expression (selective sweep) data, 245 (333) genes are annotated by GO while only 110 (145) genes are annotated by MeSH. It is important to note that this difference could be because the majority of chicken genes are annotated by Inferred from Electronic Annotation (evidence code: IEA) in GO, whereas all MeSH terms are assigned by manual curation at NCBI. On the other hand, the advantage of using GO-IEA over MeSH is that MeSH does not include genes with no published literature in PubMed, while GO-IEA can still predict function for these genes. We expect that over time, MeSH will improve as new knowledge is created and published in the scientific literature.

**Table 1:** Number of known and selected genes annotated by MeSH (Medical SubjectHeadings) and GO (Gene Ontology).

| Data | Annotated Genes |  | Selected Genes |  |  |
| --- | --- | --- | --- | --- | --- |
|  | MeSH | GO | Total | MeSH | GO |
| RNA-seq | 10227 | 12460 | 263 | 110 | 245 |
| Selective Sweep |  |  | 352 | 145 | 333 |

## Enrichment analysis

### Gene Expression Data

A subset of significant MeSH terms (*P*-value ≤ 0.05) enriched with differentially expressed genes detected in fat tissue between high and low feed efficiency chickens are highlighted in Table 2. The majority of the MeSH terms in the Chemicals and Drugs category are related to lipid deposition and lipid metabolism. For instance, *Lipoproteins* (MeSH:D008074), and *Apolipoproteins* (MeSH:D001053) are closely related to lipid transportation. Additionally, *Fatty Acid-Binding Proteins* (MeSH:D050556) regulates diverse lipid signals, while *PPAR alpha* (MeSH:D047493) controls lipid and lipoprotein metabolism. Interestingly, many GO terms related to lipid deposition and metabolism, such as *cholesterol metabolic process* (GO:0008203), *high-density lipoprotein particle assembly* (GO:0034380), *spherical high-density lipoprotein particle* (GO:0034366), and *high-density lipoprotein particle binding* (GO:0008035), were also significantly enriched with differentially expressed genes (File S1). Similarly, MeSH terms related to Wnt proteins and signalling pathways, such as *Wnt Proteins* (MeSH:D051153), *Wnt4 Protein* (MeSH: D060528), *Wntl Protein* (MeSH:D051155), and their counterparts in GO, such as *regulation of Wnt signaling pathway* (G0:0030111) and *Wnt signaling pathway* (G0:0016055), were found as significant. The Wnt proteins are known to interact with lipids. We also found *Steroid 17-alpha-Hydroxylase* (MeSH:D013254) and *steroid 17-alpha-monooxygenase activity* (G0:0004508) as significant terms; these two categories are enriched in genes involved in the synthesis of lipids. Moreover, we detected some MeSH terms related to the immune system regulation (e.g., *Interleukin-6* (MeSH:D015850) and *Chemokines* (MeSH:D018925)). Lastly, *Glycoproteins* (MeSH:D006023), is produced from the gene *AHSG* and plays a role in glucose metabolism and the regulation of insulin signaling. Taken together, our findings confirm that MeSH enrichment analysis can either reinforce findings from GO or even bring an additional biological insight. Figure 2 depicts the semantic similarity between significant MeSH terms in the Chemicals and Drugs category. In general, this subset of MeSH terms showed low to high levels of semantic similarity.

**Figure 2:**
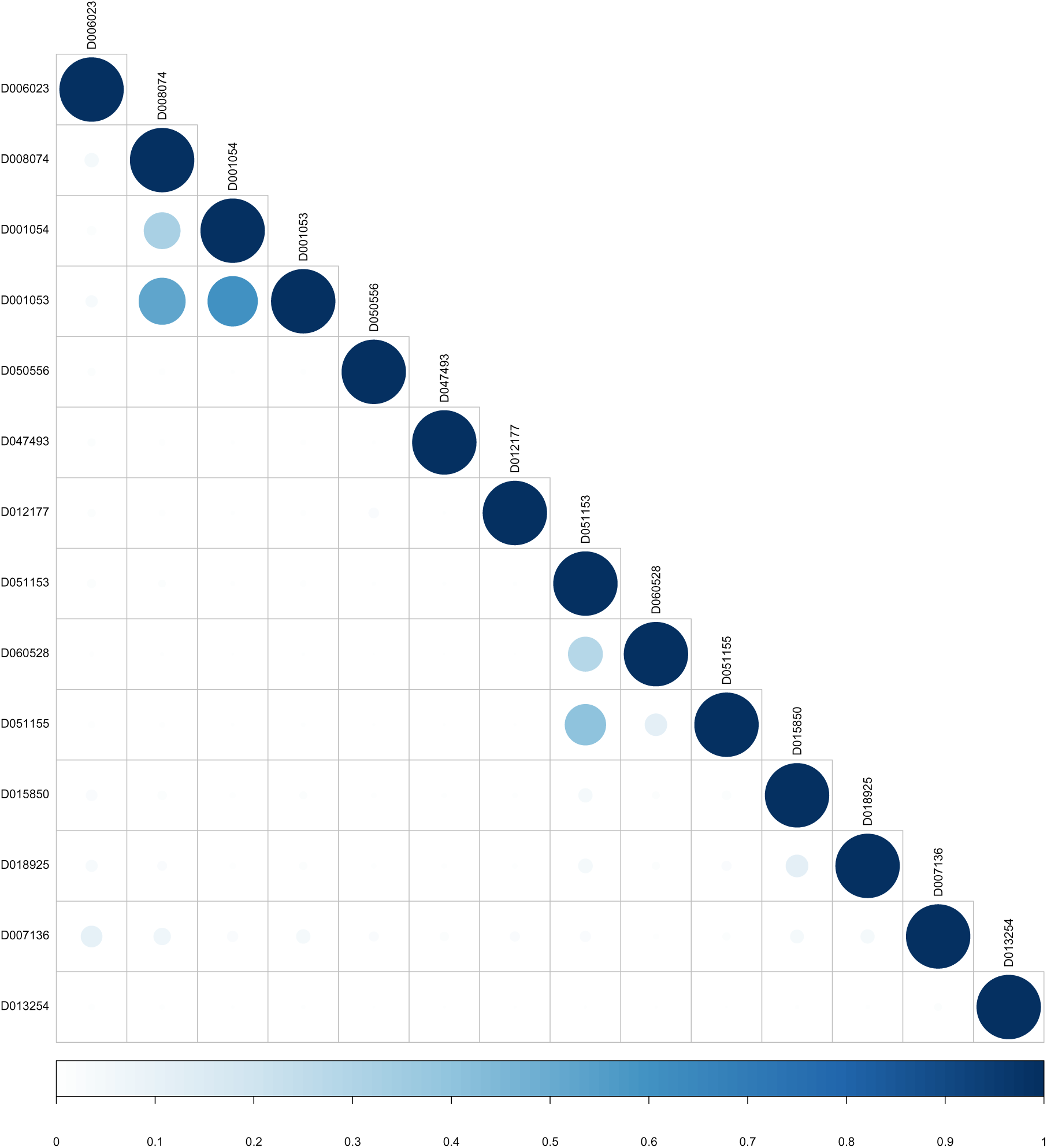
MeSH semantic similarity in the Chemicals and Drugs for the RNA-seq dataset. The higher the semantic similarity between MeSH terms, the bigger (darker) the circle. D006023 (Glycoproteins), D008074 (Lipoproteins), D001054 (Apolipoproteins A), D001053 (Apolipoproteins), D050556 (Fatty Acid-Binding Proteins), D047493 (PPAR alpha), D012177 (Retinol-Binding Proteins), D051153 (Wnt Proteins), D060528 (Wnt4 Proteins), D051155 (Wnt1 Proteins), D015850 (Interleukin-6), D018925 (Chemokines), D007136 (Immunoglobulins), and D013254 (Steroid 17alpha-Hydroxylase).

**Table 2:**
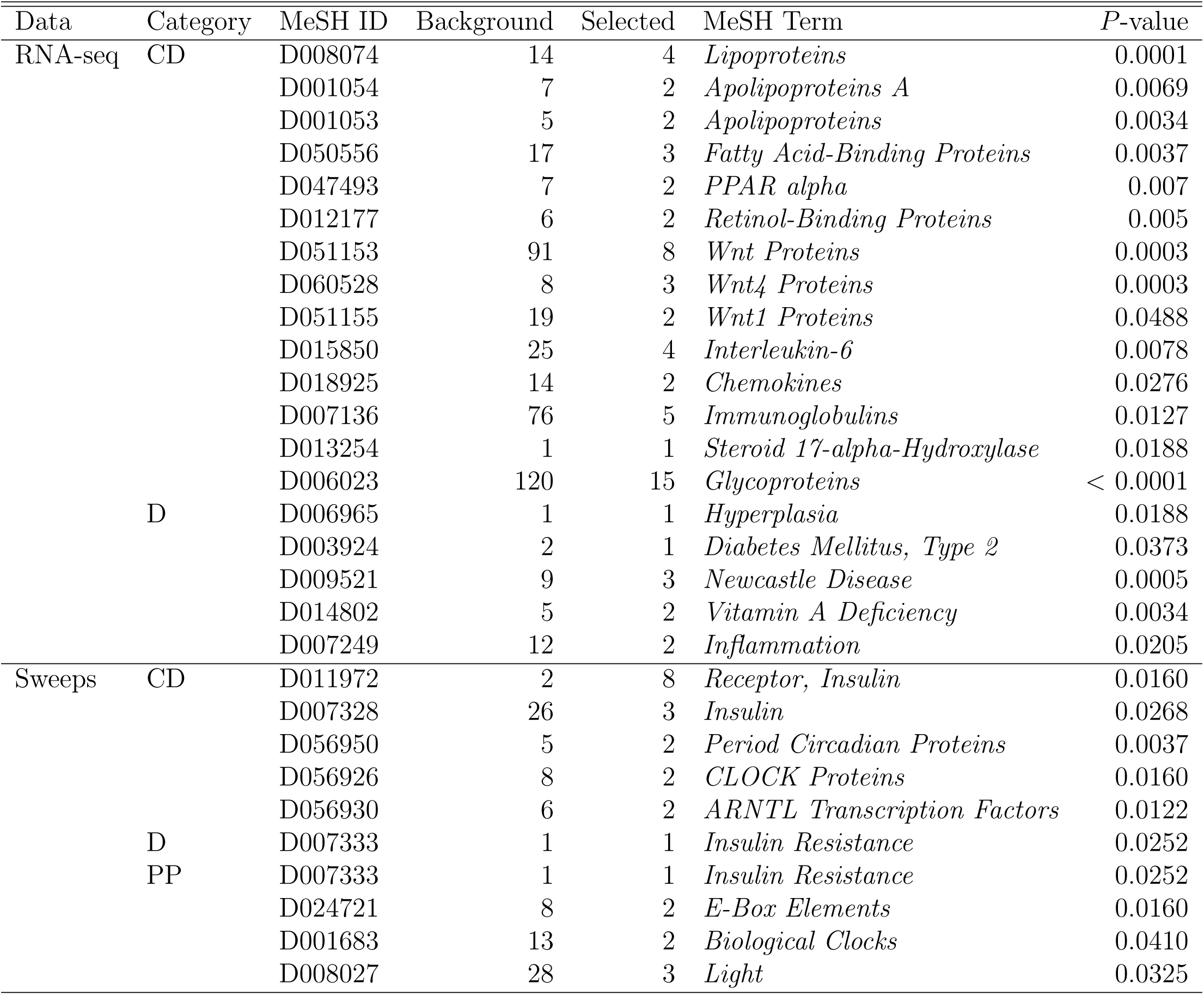
A subset of statistically signicant MeSH (Medical Subject Headings) terms. Background and Selected denote the number of background genes and selected genes annotated by the MeSH term, respectively. CD, D, and PP denote Chemicals and Drugs, Diseases, and Phenomena and Processes, respectively.

For the Diseases category, which is unique to MeSH-based analysis, a subset of significant MeSH terms that deserves particular attention in the area of feed efficiency and lipid metabolism in poultry is highlighted in Table 2. For instance, *Hyperplasia* (MeSH:D006965) is a potential contributor to abdominal fat mass in broiler chickens; its relationship with *Diabetes Mellitus, Type 2* (MeSH:D003924) is well-documented in humans. Some MeSH terms directly related to the immune function, such as *Newcastle Disease* (MeSH:D009521) and *Inflammation* (MeSH:D007249), also showed a significant enrichment with differentially expressed genes. Interestingly, *Hyperplasia* and *Inflammation* showed a moderate semantic similarity according to the MeSH hierarchy (File S1).

### Selective Sweep Data

Table 2 shows the results of the MeSH-informed enrichment analysis using genes putatively swept or under epistatic selection derived from a chicken diversity panel. Most of these terms are related to insulin metabolism. For instance, resistance to insulin occurs in birds due to high plasma glucose and fatty acid levels; this is supported by *Insulin Resistance* (MeSH:D007333) in both the Diseases and Phenomena and Processes categories, as well as *Receptor, Insulin* (MeSH:D011972) and *Insulin* (MeSH:D007328) in the Chemicals and Drugs category. Moreover, we identified MeSH terms involved in the circadian clock of chicken. These are *Period Circadian Proteins* (MeSH:D056950), *CLOCK Proteins* (MeSH:D056926) and *ARNTL Transcription Factors* (MeSH:D056930) in Chemicals and Drugs, as well as *E-Box Elements* (MeSH:D024721), *Biological Clocks* (MeSH:D001683), and *Light* (MeSH:D008027) in Phenomena and Processes. Figure 3 shows the semantic similarities among MeSH terms in the Chemicals and Drugs category. Biological clock-related annotations, such as *Period Circadian Proteins* and *CLOCK Proteins*, exhibited moderate to high similarity. The results obtained from the other MeSH and GO categories were shown in File S2.

**Figure 3:**
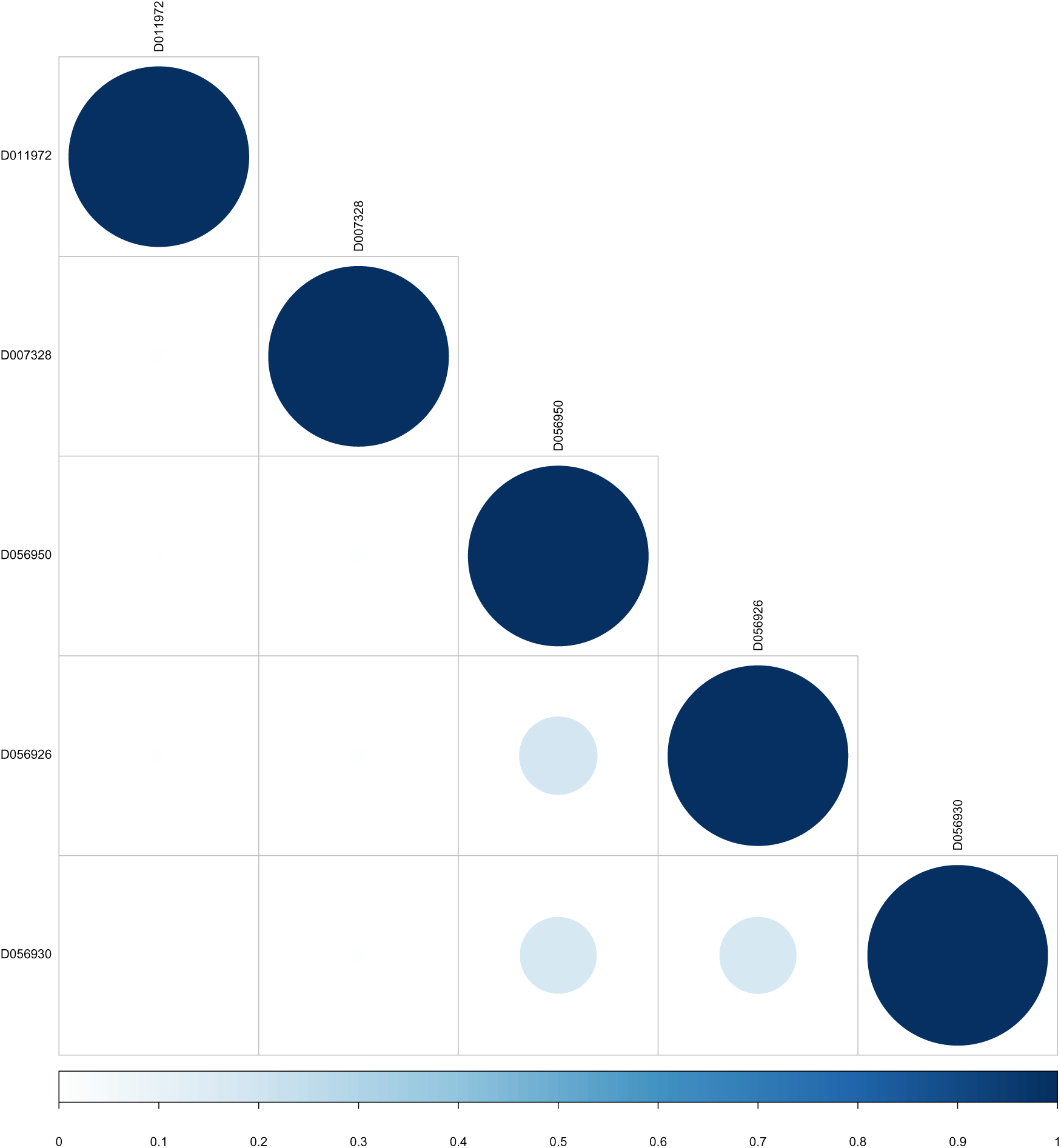
MeSH semantic similarity in the Chemicals and Drugs for the selective sweep dataset. The higher the semantic similarity between MeSH terms, the bigger (darker) the circle. D011972 (Receptor, Insulin), D007328 (Insulin), D056950 (Period Circadian Proteins), D056926 (CLOCK Proteins), and D056930 (ARNTL Transcription Factors).

### Gene semantic similarity

#### Gene Expression Data

Comparison of gene semantic similarity between MeSH and GO Biological Process for a subset of significant genes (*n*=49) from the RNA-seq dataset is depicted in Figure 4. MeSH-based gene semantic similarity analysis showed that genes related to energy reserve metabolic process are highly related. For instance, genes that are involved in triacylglycerol and cholesterol biosynthesis, such as methylsterol monooxygenase 1 (*MSMO1*), insulin induced gene 1 (*INSIG1*), 1-acylglycerol-3-phosphate O-acyltransferase 9 (*AGPAT9*), and ADP ribosylation factor like GTPase 2 binding protein (*ARL2BP*), were highly similar to each other based on the MeSH hierarchy. Interestingly, GO-based analysis produced slightly different results; for instance, the gene *MSMO1* was highly similar to *INSIG1* but moderately similar to *AGPAT9* and *ARL2BP*. Additionally, genes *MSMO1* and *INSIG1* were moderately or highly related to lecithin-cholesterol acyltransferase (*LCAT*) and cytochrome b5 type A (microsomal) (*CYB5A*) based on the GO structure. These two genes, involved in lipid metabolism, also showed high similarity to apolipoprotein A-I (APOA1) and cytochrome P450, family 17, subfamily A, polypeptide 1 (*CYP17A1*). The relationship among these genes were low to moderate based on the MeSH hierarchy. The results based on the GO Molecular Function and Cellular Component categories were presented in File S3.

**Figure 4:**
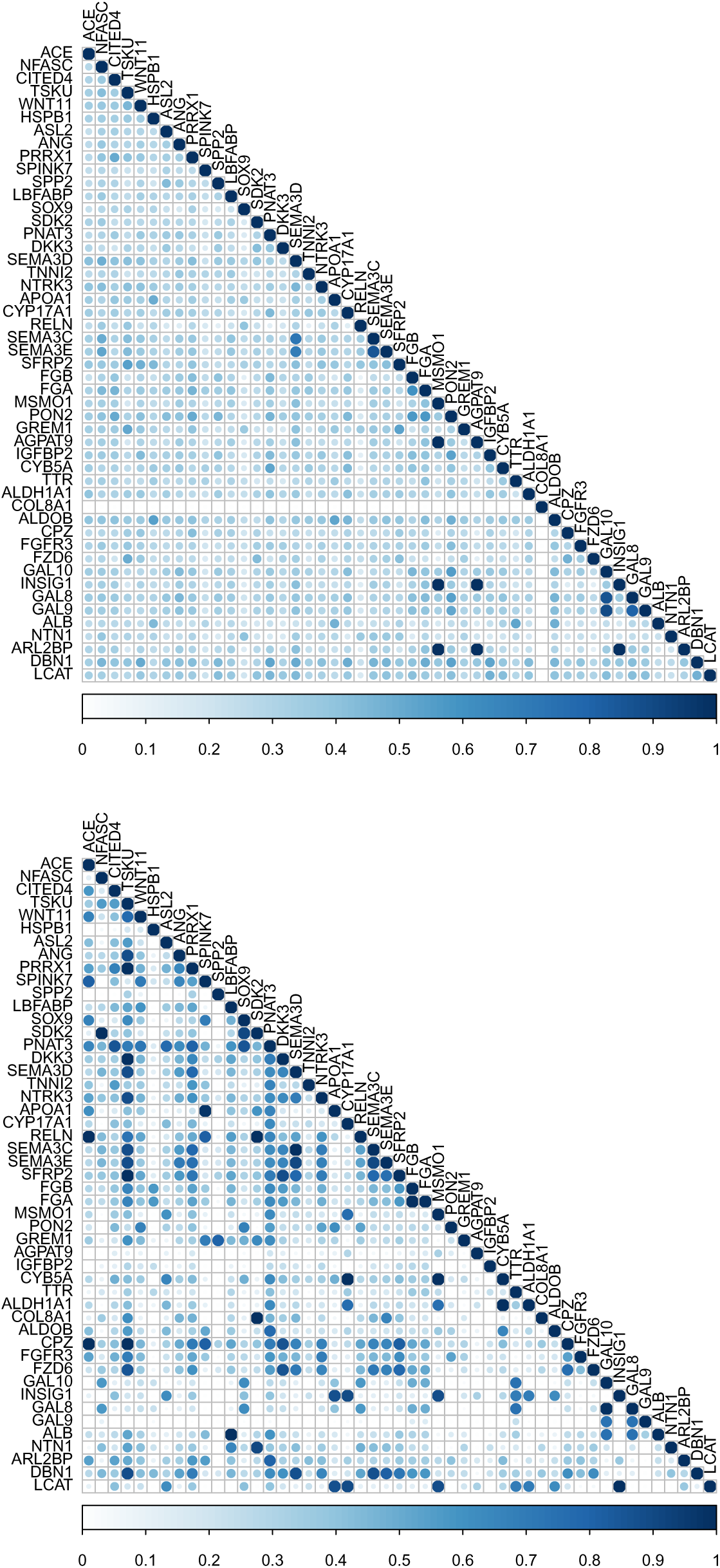
Gene semantic similarity for the RNA-seq dataset. The higher the semantic similarity between gene pairs, the bigger (darker) the circle. Top:MeSH, Bottom:GO.

#### Selective Sweep Data

Gene semantic similarity based on both MeSH and GO Biological Process among a subset of genes (*n*=45) under selection is shown in Figure 5. Notably, a large group of genes, including strawberry notch homolog 1 (Drosophila) (*SBNO1*), ARP5 actin-related protein 5 (*ACTR5*), SET domain containing 1B (*SETD1B*), Obg-like ATPase 1 (*OLA1*), and histone deacetylase 9 (*HDAC9*) were highly related based on both MeSH and GO-guided semantic similarity analyses. All these genes are involved in chromatin organization and regulation of gene expression. Moreover, particular attention was paid to the top five candidates under epistatic selection reported by Beissinger et al. (2015). These genes are adenylate cyclase 5 (ADCY5), myosin light chain kinase (*MYLK*), phosphatidylinositol-4,5-bisphosphate 3-kinase, catalytic subunit beta (*PIK3CB*), calcium binding protein 39 (*CAG39*), and interleukin 1 receptor accessory protein (*IL1RAP)*. Although none of these pair of genes appeared in a GO-based similarity matrix, *ADCY5* and *MYLK* presented a low to moderate gene semantic similarity based on the MeSH hierarchy (File S4).

**Figure 5:**
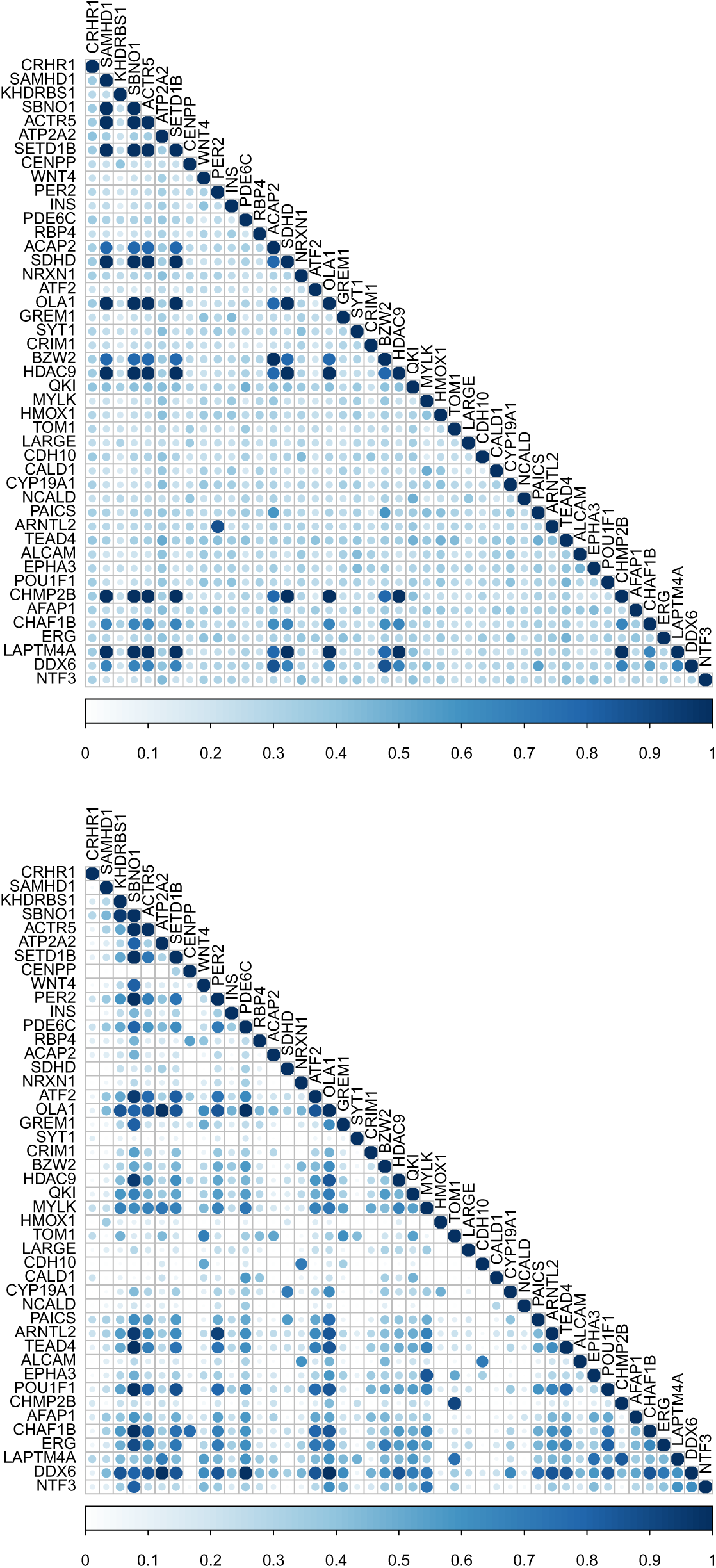
Gene semantic similarity for the selective sweep dataset. The higher the semantic similarity between gene pairs, the bigger (darker) the circle. Top:MeSH, Bottom:GO.

## Discussion

This article reports the MeSH analysis for chicken using the newly developed Bioconductor packages. These new resources enabled us to carry out different MeSH-based analyses, including enrichment analysis and MeSH-guided semantic similarity among functional terms and gene products. We exemplified the potential usefulness of these MeSH-based approaches by using two different publicly available chicken data.

The adipose tissue is the major site for lipid deposition and lipid metabolism, and it plays a central role in energy homeostasis. Unsurprisingly, several MeSH terms closely related to fat metabolism, such as Lipoproteins, Apolipoproteins, Fatty Acid-Binding Proteins, and PPAR alpha, were significantly enriched with genes that showed differential expression in fat tissue between high and low feed efficiency broiler chickens. We found some genes were annotated by the same MeSH terms. For instance, gene overlap between Lipoproteins and Apolipoproteins was one-half and 66% of genes were shared between Fatty Acid-Binding Proteins and PPAR alpha. It is likely that this gene overlap is observed because each MeSH term inherits all annotations from its more specific child terms (Falcon and Gentleman, 2007). It is possible to address this issue by conducting a conditional analysis that is implemented in the GOstats package. Adding this feature in the meshr package might alleviate the overlap of genes. Also, adipose tissue is now recognized as a metabolically active tissue that has important endocrine and immune regulatory functions (Kershaw and Flier, 2004). Interestingly, we found many significant MeSH terms, such as Interleukin-6, Chemokines, and Immunoglobulins, that are closely associated with the regulation of the immune function. Overall, our MeSH-based findings provide further insights into the biological mechanisms underlying differences in adiposity between high and low feed efficiency broiler chickens.

Included in our exemplary applications of MeSH annotations is a set of 352 genes previously identified as putatively affected by selection. Genes identified through population-genetic approaches such as this can be elusive, because their identification does not rely on phenotypes. Therefore associating selection with any specific trait is often very difficult (Akey, 2009). As we demonstrate in this study, tools such as GO and now MeSH are useful for suggesting biological interpretations that can later be followed up on or drive future biological hypotheses. For instance, our results showed that insulin-related MeSH terms appeared unusually often in the set of genes impacted by selection. This implies that selection for insulin-related traits may have played an important role in differentiating chicken breeds. Furthermore, our analysis involved testing for semantic similarity between pairs of genes, which was particularly useful for evaluating the most promising gene-pairs highlighted by Beissinger et al. (2015) as candidates for epistatic selection. Our expectation was that these pairs of genes are likely to be related to each other, as they have been predicted to be involved in the same selected phenotype. Our finding that one pair showed at least a weak semantic similarity may be interpreted as evidence that these two genes, *ADCY5* and *MYLK* are the most likely among the set to truly be epistatic.

The recent advancement in cataloguing genes with MeSH and GO has made it possible to assess the role of selected genes and has opened new opportunities for genetic research. Enrichment analysis recapitulates a set of genes into higher-level biological features. We argue that obtaining a complete picture of genes of interest using MeSH and GO is an important initial step toward functional genomics studies in poultry as well as other agricultural species as it facilitates efforts to illuminate the genetic basis of phenotypic variation.

## Acknowledgements

This work was supported in part by the University of Nebraska Layman Fund to G.M. Support for T.M.B. was provided by USDA-ARS. F.P. acknowledges financial support from the Florida Agricultural Experiment Station and the Department of Animal Sciences, University of Florida.

## Supporting Information

- File S1: MeSH over-representation analysis (RNA-seq data)
- File S2: MeSH over-representation analysis (Selective sweep data)
- File S3: Gene Semantic Similarity (RNA-seq data)
- File S4: Gene Semantic Similarity (Selective sweep data)

